# Interactive effects of temperature, cadmium, and hypoxia on rainbow trout *(Oncorhynchus mykiss)* liver mitochondrial bioenergetics

**DOI:** 10.1101/2024.07.15.603625

**Authors:** John O. Onukwufor, Collins Kamunde

**Author notes:** **Corresponding Authors**: Department of Biomedical Sciences, Atlantic Veterinary College, University of Prince Edward Island, 550 University Avenue, Charlottetown, PE, Canada C1A 4P3.

## Abstract

Fish in their natural environments possess elaborate mechanisms that regulate physiological function to mitigate the adverse effects of multiple environmental stressors such as temperature, metals, and hypoxia. We investigated how warm acclimation affects mitochondrial responses to Cd, hypoxia, and acute temperature shifts (heat shock and cold snap) in rainbow trout. We observed that state 3 respiration driven by complex I (CI) was resistant to the stressors while warm acclimation and Cd reduced complex I +II (CI + II) driven state 3 respiration. In contrast, state 4 (leak) respirations for both CI and CI + II were consistently stimulated by warm acclimation resulting in reduced mitochondrial coupling efficiency (respiratory control ratio, RCR). Warm acclimation and Cd exacerbated their individual effect on leak respiration to further reduce the RCR. Moreover, the effect of warm acclimation on mitochondrial bioenergetics aligned with its inhibitory effect on activities of citrate synthase and both CI and CII. Unlike the Cd and warm acclimation combined exposure, hypoxia alone and in combination with warm acclimation and/or Cd abolished the stimulation of CI and CI + II powered leak respirations resulting in partial recovery of RCR. The response to acute temperature shifts indicated that while state 3 respiration returned to pre-acclimation level, the leak respiration did not. Overall, our findings suggest a complex *in vivo* interaction of multiple stressors on mitochondrial function that are not adequately predicted by their individual effects.

**Highlights:**

- Mitochondrial bioenergetics plasticity was investigated.
- Stimulation of mitochondrial leak state is a key effect of warm acclimation.
- State 3 but not leak respiration returns to control status after warm acclimation.
- Warm acclimation and Cd additively stimulate leak respiration and reduce RCR.
- Hypoxia protects against deleterious effects of warm acclimation and Cd exposure.

**Graphical abstract:** 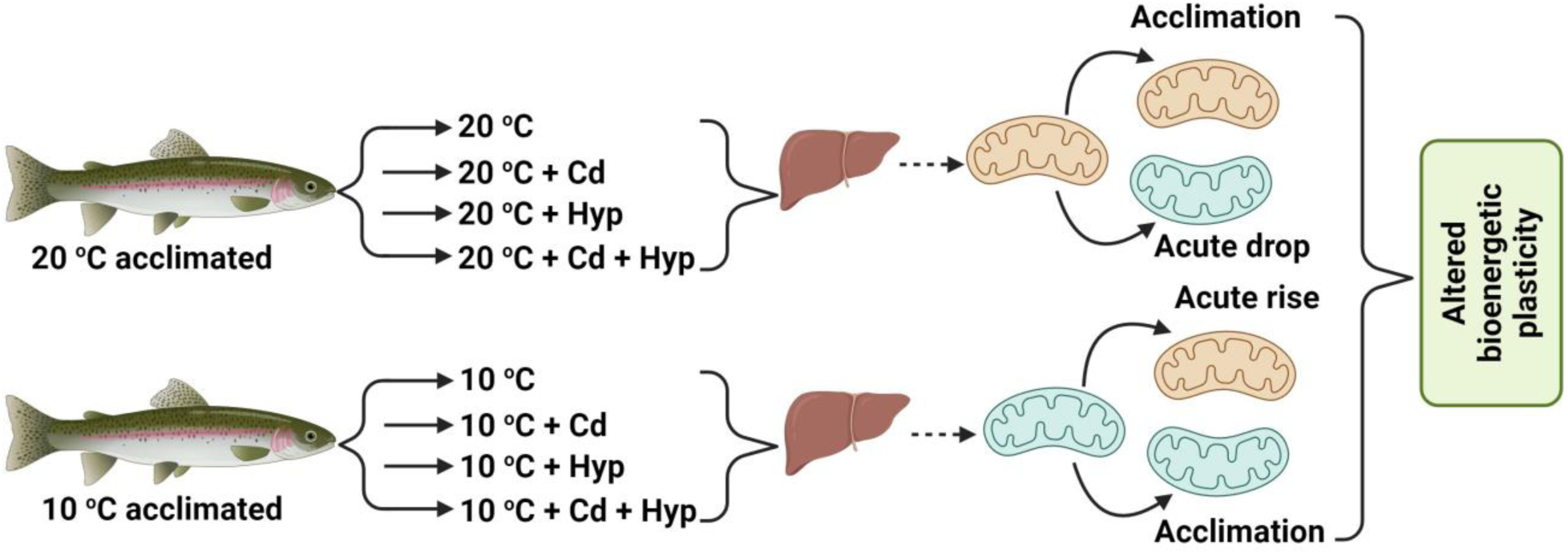

## 1. Introduction

A wide array of environmental stressors impairs mitochondrial function (Belyaeva and Korotkov, 2003; Sokolova, 2004; Kurochkin et al., 2009; Adiele, et al., 2010; 2012; Ivanina et al., 2012; Sappal et al., 2014; 2015; Onukwufor et al., 2014; 2015; 2016; 2017). Because of their central role in multiple critical cellular processes (Naquet et al., 2016; Bohovych and Khalimonchuk, 2016; Vakifahmetoghu-Norberge et al., 2017), impairment of mitochondrial functional integrity has implications for cellular survival and organismal health. While the exact mechanisms by which different stressors impair mitochondrial functions are diverse, altered electron transport system (ETS) activity is a common sequel of exposure to multiple stressors (Lee et al., 2005; Orlvo et al., 2013; Onukwufor et al., 2014; 2015; 2016; 2017). In this regard, *in vitro* studies using isolated mitochondria are important in identifying the mechanisms of action of stressors, but they are limited in elucidating adaptive responses of mitochondria to environmental stress. Aquatic organisms have been shown to adapt their mitochondria following long term exposure to environmental stressors (Hazel and Prosser, 1974; Hochacka and Somero, 1984). These adaptive responses are geared toward re-establishing homeostasis during chronic stress and include changes in membrane phospholipids, enzyme activity, substrate affinity and preference, and antioxidant defense system capacity (Hazel, 1995; Guderley and St-Pierre, 2002; Guderley, 2004; Kraffe et al., 2007; Lockwood and Somero, 2012; Oellermann, et al., 2012).

Temperature fluctuations due to natural phenomena or as a consequence of global climate change (Portner, 2002; Portner and Knust, 2007; Portner and Farrell, 2008; Somero, 2010) and hypoxia as a result of eutrophication and diurnal/seasonal changes in O_2_ availability in water (Wu, 2002; Hattink et al., 2005; Diaz and Rosenberg, 2008) are two highly consequential stressors in the aquatic environment. Temperature stress and hypoxia can occur together or sequentially leading to alterations in how fish response to stress induced by metals such as cadmium (Cd) (Onukwufor et al., 2017). Notably, it has been shown that temperature, hypoxia, and Cd target the mitochondria (Sokolova, 2004; Kurochkin et al., 2009; Onukwufor et al., 2017). Based on *in vitro* studies, combinations of these stressors impair mitochondrial bioenergetics to a greater extent compared with the respective stressors acting individually (Sappal et al., 2014; 2015; Onukwufor et al., 2014; 2015; 2016; 2017). However, it remains to be investigated whether these *in vitro* mitochondrial responses occur *in vivo* wherein exposed organisms can mount adaptive mechanisms to counteract effect of stressors.

In the present study, we probed how warm acclimation affects mitochondrial responses to subsequent acute temperature shifts and exposure to Cd and hypoxia in rainbow trout *(Oncorhynchus mykiss) in vivo*. First, we tested the idea that effects of *in vivo* exposure to temperature, hypoxia, and Cd would be similar to those observed in *in vitro* studies (Onukwufor et al., 2014; 2015; 2016; 2017). We predicted that effects of these stressors would be exacerbated when acting in binary or ternary combinations compared with their individual effects. Second, we hypothesized that liver mitochondrial respiratory function in fish acclimated to warm temperature would be similar to that of un-acclimated fish. Third, we hypothesized that warm acclimation would temper mitochondrial responses to acute thermal shifts and exposure to hypoxia and Cd.

## 2. Materials and Methods

### 2.1. Ethics

The study and all the experimental procedures were approved by the University of Prince Edward Island Animal Care Committee in accordance with the Canadian Council on Animal Care (protocol # 17-036).

### 2.1. Fish and temperature acclimation

Juvenile female diploid rainbow trout weighing 200 to 400 g were purchased from Ocean Farms Inc., Brookvale, PE, and maintained in the Aquatic Facility of the Atlantic Veterinary College in a 400-l tank for 2 weeks at 10 ^°^C. The fish were then divided into two groups earmarked for acclimation to 10 ^°^C (control group) and 20 ^°^C (warm-acclimated group). For warm acclimation, the water temperature was raised from 10 ^°^C by 1 ^°^C/day to 20 ^°^C over 10 days. The fish were acclimated to the two temperature regimes for 50 days and thereafter exposed to two dissolved oxygen (DO) levels (> 90% and 30% air saturation) and two Cd concentrations (0 and 10 µg/l) alone and in combination.

### 2.2. Experimental protocol for Cd exposure and hypoxia challenge

Fish were fasted for 24 h and moved from the holding tanks to an experimental room maintained at 20 ^°^C. Two 400-l tray holding tanks were used as water baths with one maintained at 10 and the other at 20 ^°^C. The temperature of the 10 ^°^C water bath was regulated with a chiller while heating/cooling was not required for the 20 ^°^C water bath because it was equivalent to room temperature. The two tanks each held two 40-l experimental containers with lids. A total of 8 fish were sampled on each sampling day, 4 fish each from the 10 and 20 ^°^C holding tanks. Two fish were placed in the 40-l tanks under constant aeration and allowed to settle for 30 min. For Cd exposure, 10 µg/l of Cd was added, and the exposure was done for 24 h. Hypoxic conditions were generated by bubbling N_2_ into the 40-l tank to reduce the DO level in the water to 30% air saturation (took 2-5 min). The hypoxic conditions were maintained by balancing the bubbling of N_2_ and air with constant monitoring of the DO level with an oxygen probe (YSI model 55, Yellow Spring, Ohio, USA). The fish were then exposed to the hypoxic conditions for 2 h. For combined hypoxia and Cd exposure, fish were first exposed to 10 µg/l Cd for the first 22 h following which N_2_ was bubbled to deplete DO to 30% and maintained over the last 2 h of the 24 h Cd-exposure. Thus, in total, both the warm- and 10 ^°^C-acclimated fish were exposed to Cd and hypoxia for 24 and 2 h, respectively. The measured Cd and DO levels were within 5% of the nominal levels.

### 2.3. Mitochondrial isolation

At the end of the *in vivo* Cd and hypoxia exposures, fish were stunned with a blow to the head, decapitated, and the livers were immediately harvested. Mitochondrial isolation was done as described (Onukwufor et al., 2014). Briefly, each liver was rinsed with mitochondrial isolation buffer (MIB: 250 mM sucrose, 10 mM Tris-HCl, 10 mM KH_2_PO_4_, 0.5 mM EGTA, 1 mg/ml BSA (fatty acid free), 2 µg/ml aprotinin, pH 7.3), blotted dry, and weighed. Thereafter, each liver was diced and homogenized in 1:3 (weight to volume) ratio of liver to MIB in a 10-ml Potter-Elvehjem homogenizer (Cole Parmer, Anjou, QC). Three passes of the pestle mounted on a hand-held drill (MAS 2BB, Mastercraft Canada, Toronto, ON) running at 200 rpm were optimal for rainbow trout liver homogenisation. The homogenate was initially centrifuged at 800 ×*g* for 15 min at 4 ^°^C. The supernatant was collected, centrifuged at 13,000 ×*g* for 10 min at 4 ^°^C and the resulting mitochondrial pellet was washed twice by re-suspending in MIB and centrifuging at 11,000 ×*g* for 10 min at 4 ^°^C. The mitochondrial pellet from the last wash was re-suspended in a 1:3 (weight to volume) ratio of mitochondrial respiration buffer (MRB: 10 mM Tris-HCl, 25 mM KH_2_PO_4_, 100 mM KCl, 1 mg/ml BSA (fatty acid free), 2 µg/ml aprotinin, pH 7.3). The suspension was divided into two portions, one for mitochondrial respiration measurement and the other for measurement of activities of mitochondrial enzymes. The latter was stored at -80 ^°^C and the enzyme analyses were done within 2 weeks.

### 2.4. Mitochondrial respiration

The protein concentrations of mitochondrial subsamples earmarked for respiration measurements were determined by spectrophotometry (Spectramax Plus 384, Molecular Device, Sunyvale, CA) according to Bradford (1976). Mitochondrial respiration was then measured at the fish acclimation temperatures of 10 and 20 ^°^C except for the determination of thermal plasticity of mitochondrial function where 10 ^°^C-acclimated fish were assessed at 20 ^°^C and warm-acclimated fish were assessed at 10 ^°^C. The assay temperatures were maintained using a recirculating water-bath (Haake, Karlsruhe, Germany). Respiration measurements were done using Clark-type oxygen electrodes (Qubit System, Kingston, ON) in 1.5-ml assay volume after a two-point calibration at 0 and 100% air saturation at the ambient atmospheric pressure. At the end of the calibration, 1.45 ml of MRB and 100 µl of mitochondrial suspension (2.3-2.8 mg of protein) were loaded into the continuously stirred cuvettes. First, 5 mM each of glutamate and malate (glutamate-malate) were added followed by 0.25 mM ADP to impose complex I (CI) state 3 respiration (oxidative phosphorylation state, OXPHOS state) which transitioned to state 4 (leak state) respiration on complete phosphorylation of the added ADP. Complex II (CII) substrate, succinate (I mM), was then added resulting in convergent electron flux wherein electrons from CI and CII are simultaneously fed to the Q-pool. Here, addition of the second bolus (0.25 mM) of ADP induced CI + CII state 3 respiration which transitioned to state 4 respiration upon complete phosphorylation of the added ADP. Thereafter, 2.5 µg/ml of oligomycin was added to inhibit ATP synthase, allowing the determination of state 4+ respiration, an estimate of mitochondrial leak (Brand et al., 1994; St-Pierre et al., 2000). The respiratory control ratio (RCR(+)) was subsequently calculated from states 3 and state 4(4+) rates of respiration rates according to Chance and Williams (1955). Note that except for the acute temperature shift (10 ⇔ 20 ^°^C), interactive effects of the stressors on mitochondrial respiration were measured using freshly isolated organelles following *in vivo* exposure to the stressors without additional *in vitro* exposure to the stressors.

### 2.5. Activities of mitochondrial enzymes

Mitochondrial samples stored at -80 ^°^C were thawed and equal volumes of each sample and 2% Triton X-100 were mixed, sonicated on ice for 10 s and the protein concentrations were measured. The samples were then used to measure activities of citrate synthase and CI and CII.

Citrate synthase activity was measured as previously described (Onukwufor et al., 2014). Briefly, an assay buffer containing 1 M/l Tris-HCl, 2 mM 5,5’-dithiobis-(2-nitrobenzoic acid) (DTNB), 2 mM Acetyl coenzyme A and 1 % (v/v) Triton X-100 was constituted. Samples were run in duplicates in 96-well microplates by adding 50 µl of the assay reagent, 40 µl (40 µg) mitochondrial suspension and 150 µl of Millipore water. To start the reaction, 10 µl of 12.5 mM oxaloacetic acid was added to the plate. In this assay, citrate synthase catalyses the reaction of oxaloacetic acid with acetyl-CoA to form citrate and CoA-SH. The CoA-SH then reacts with DTNB to form TNB which has an intense yellow color at 412 nm. The samples were run with and without oxaloacetic, and absorbance at 412 nm was read every 15 s for 10 min. Citrate synthase activity was calculated from the increase in absorbance by subtracting the values obtained without added oxaloacetic acid from the values obtained with added oxaloacetic acid. The specific enzyme activity was expressed in molar units using a molar extinction coefficient of 13.6 mmol^-1^ cm^-1^ for TNB.

For CI activity, 240 µl of assay buffer (25 mM potassium phosphate, 3.5 mg/ml BSA, 100 µM 2,6-dichlorophenol indophenol (DCIP), 70 µM decylubiquinone, 0.6 mg/l antimycin A, and 200 µM NADH, pH 7.3) were added to each microplate well. To start the reaction, 60 µg (in 10 µl) of mitochondrial protein were added to all wells except the blanks (10 µl of assay buffer were added) and each sample was analyzed in triplicate with and without 2 µM rotenone. The decrease in absorbance due to the reduction of DCIP, the terminal electron acceptor in this assay, was monitored spectrophotometrically (Spectramax 384 Plus) at 600 nm for 5 min at 15 s intervals. CI activity was then calculated by subtracting the rotenone-insensitive activity from the total activity and was converted to micromoles of DCIP reduced using a molar extinction coefficient of 19.1 mM^-1^ cm^-1^.

To measure CII activity, an assay buffer containing 250 mM potassium phosphate, 50 mM MgCl_2_, 10 mM DCIP, 1 M KCN, 1 mM rotenone, 1 mg/l antimycin A and 1 M sodium succinate, pH 7.3 was constituted. Thereafter, 34 µl of the assay buffer and 10 µl (60 µg) of mitochondrial suspension were added to a 96 well microplate in triplicate and the volume brought to 240 µl by adding Millipore water. To establish the baseline, absorbance was read at 600 nm for 3 min. The addition of 10 µl of 1.625 mM of coenzyme Q_1_ initiated the reaction. The resulting decrease in absorbance due to the reduction of DCIP was recorded every 15 s for 10 min at 600 nm (Spectramax 384 Plus). A molar extinction coefficient of 19.1 mM^-1^ cm^-1^ for DCIP was used to convert CII activity to molar units.

### 2.6. Statistical analysis

Statistical analysis was done using Sigmaplot 11.0 (Systat Software, San Jose, CA, USA). All the data were first tested for normality and equality of variances before running analysis of variance (ANOVA). Data that failed ANOVA assumptions were subjected to square root transformation. Data that passed the tests after transformation were submitted to ANOVA and those that still failed were analyzed using Kruskal-Wallis ANOVA on ranks. Data were analyzed by 3-way ANOVA with hypoxia, temperature, and Cd as the independent variables. Significantly different means were separated using Tukey’s HSD *post hoc* test at p < 0.05.

## 3.0. Results

### 3.1. Effects and interactions of temperature, Cd, and hypoxia on CI respiration

Individually, hypoxia (F_1,32_ = 0.002; p = 0.97), temperature (F_1,32_ = 1.380; p = 0.25), and Cd (F_1,32_ = 1.28; p = 0.27) did not alter CI state 3 respiration (Fig. 1A). However, the temperature × Cd interaction was significant (F_1,32_ = 4.74; p = 0.037) indicating that the effect of Cd depended on the temperature. CI state 4 respiration (Fig. 1B) was significantly altered by hypoxia (F_1,32_ = 7.07; p = 0.012) and temperature (F_1,32_ = 18.5; p < 0.001) but not Cd (F_1,32_ = 1.73; p = 0.198). Here, 10 °C-acclimation singly and jointly with hypoxia resulted in the lowest CI state 4 respiration rates while warm-acclimation singly and in combination with Cd evoked the highest rates. Complex I RCR (Fig. 1C) was altered by hypoxia (F_1,32_ = 29.9; p < 0.001), temperature (F_1,32_ = 134.8; p < 0.001), and Cd (F_1,32_ = 34.15; p < 0.001) with significant interactions of hypoxia × Cd (F_1,32_ = 7.5; p = 0.01) and temperature × Cd (F_1,32_ = 65.14; p < 0.001). Aligned with the effects of the treatments on CI state 3 (Fig. 1A) and 4 respiration rates (Fig. 1B), warm-acclimated groups had lower RCR (Fig. 1C) than their 10 °C-acclimated counterparts. Notably, the RCR of mitochondria of the 10 °C-acclimated fish was 5-fold higher than the warm-acclimated and was reduced by all treatments except hypoxia.

**Figure 1:**
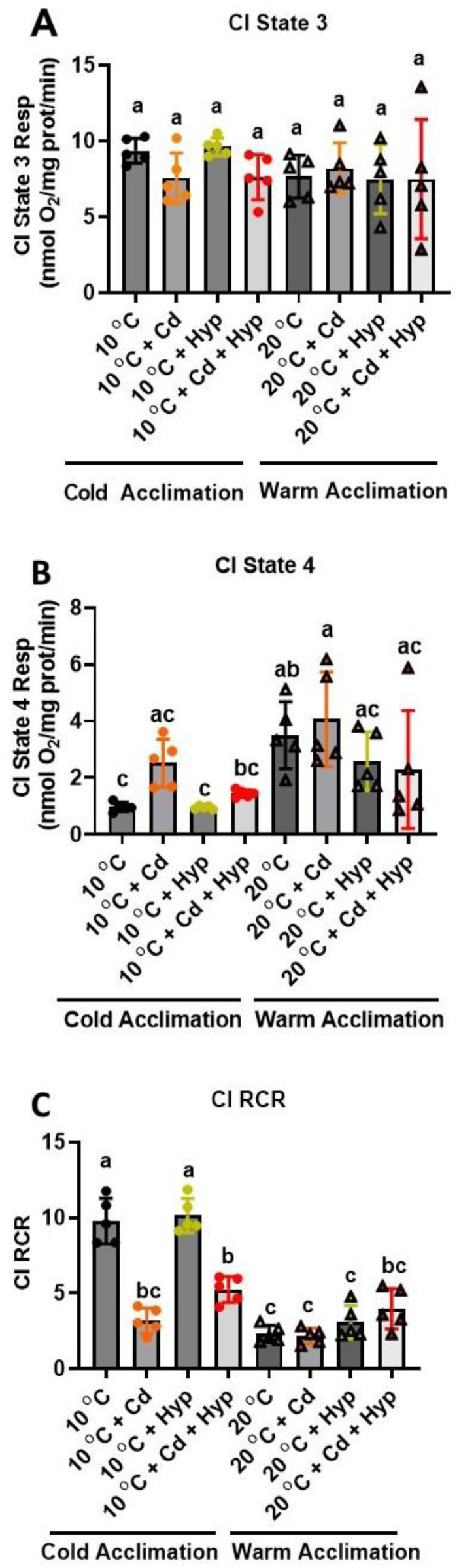
Individual and combined effects of acclimation temperature, Cd, and hypoxia on mitochondrial complex I powered respiration. (**A**) state 3, (**B**) state 4, and (**C**) RCR. Rainbow trout were acclimated to 10 °C (control) or 20 °C (warm-acclimated) for 50 days and exposed to (i) 10 µg/l Cd for 24 h, (ii) hypoxia (30% air saturation) for 2h, or (iii) 10 µg/l Cd for 24 h combined with hypoxia (30% air saturation) for 2 h. Liver mitochondria were isolated and the respiration fueled by glutamate-malate was measured at the respective acclimation temperature. Data are means ± SEM, N = 5 independent fish. Bars with different letters represent statistically significant means (p < 0.05), three-way ANOVA, Tukey’s HSD test.

### 3.2. Effects and interactions of temperature, Cd, and hypoxia on CI + CII respiration

To unveil the effect of the stressors on maximum mitochondrial respiratory capacity, we exploited the principle of convergent electron flow from CI and CII to the Q-junction by energizing mitochondria with glutamate+malate+succinate. We found that CI + CII state 3 respiration rate (Fig. 2A) in control (10 ^°^C-acclimated fish not exposed to Cd and/or hypoxia) was higher than that for CI alone (Fig. 1A vs. Fig. 2A). Furthermore, the CI + II state 3 respiration rate was not altered by hypoxia (F_1,32_ = 1.01; p = 0.32) but was reduced by warm acclimation (F_1,32_ = 27.08; p < 0.001) and Cd exposure (F_1,32_ = 4.56; p = 0.04) (Fig. 2A). CI + CII state 4 respiration rate was significantly altered by hypoxia (F_1,32_ = 14.0; p < 0.001) and temperature (F_1,32_ = 12.18; p = 0.001) but not Cd (F_1,32_ = 2.70; p = 0.11) (Fig. 2B), and the interactions were not significant. Specifically, warm-acclimation increased state 4 respiration rate while hypoxia alone and in combination with Cd blunted this increase. The response pattern of CI + CII state 4+ respiration (Fig. S1A) mirrored that of state 4; however, only the effects of hypoxia (F_1,32_ = 8.72; p = 0.006) and the hypoxia × temperature interaction (F_1,32_ = 4.66; p = 0.039) were significant.

**Figure 2:**
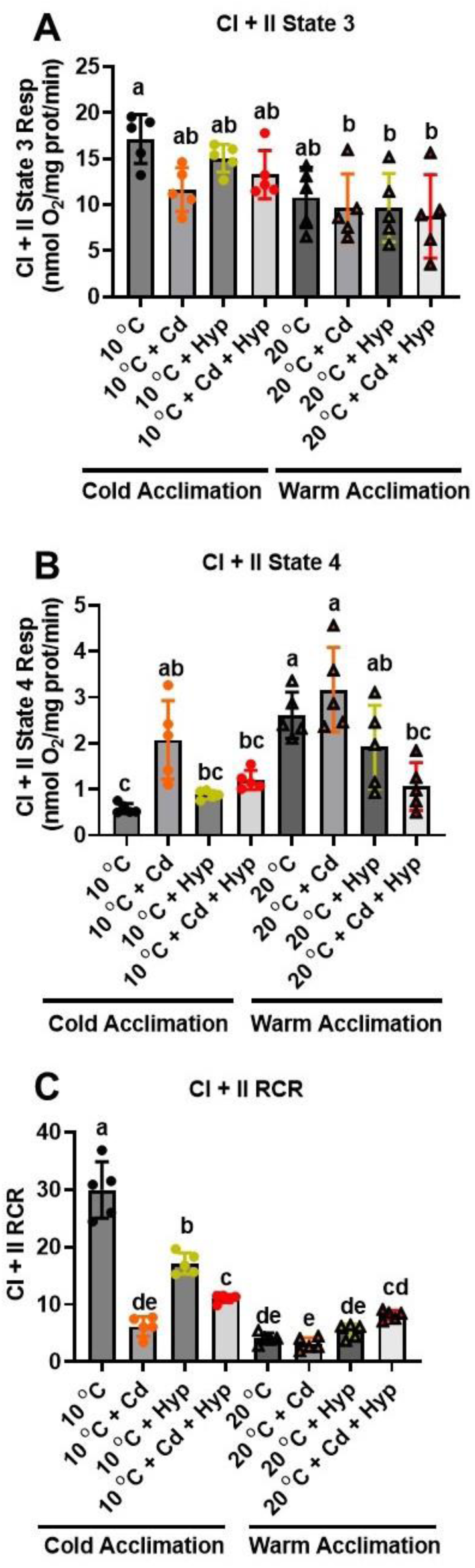
Individual and combined effects of acclimation temperature, Cd, and hypoxia on mitochondrial complex I + II powered respiration. (**A**) state 3, (**B**) state 4, and (**C**) RCR. Rainbow trout were acclimated to 10 °C (control) or 20 °C (warm-acclimated) for 50 days and exposed to (i) 10 µg/l Cd for 24 h, (ii) hypoxia (30% air saturation) for 2h, or (iii) 10 µg/l Cd for 24 h combined with hypoxia (30% air saturation) for 2 h. Liver mitochondria were isolated and the respiration fueled by glutamate-malate-succinate was measured at the respective acclimation temperature. Data are means ± SEM, N = 5 independent fish. Bars with different letters represent statistically significant means (p < 0.05), three-way ANOVA, Tukey’s HSD test.

Consistent with the observed changes in state 3 and 4 respiration rates, CI + CII RCR was reduced by warm temperature acclimation (F_1,32_ = 152.6; p < 0.001), Cd (F_1,32_ = 21.2; p < 0.001) and hypoxia (F_1,32_ = 33.96; p < 0.001). Additionally, the interactions of temperature × hypoxia (F_1,32_ = 7.77; p = 0.009), temperature × Cd (F_1,32_ = 47.81; p = 0.001) and hypoxia × Cd (F_1,32_ = 24.48; p = 0.001) were significant (Fig. 2C). In contrast, the RCR^+^ for CI + CII was not affected by hypoxia (F_1,32_ = 2.91; p = 0.10) but was reduced by warm acclimation (F_1,32_ = 73.26; p < 0.001) and Cd (F_1,32_ = 19.2; p < 0.001) (Fig. S1B). Furthermore, the interactions of hypoxia × temperature (F_1,32_ = 4.38; p = 0.04), hypoxia × Cd (F_1,32_ = 16.66; p < 0.001), and temperature × Cd (F_1,32_ = 29.92; p < 0.001) were significant.

### 3.3. Effects of hypoxia, temperature, and Cd on activities of mitochondrial enzymes

Warm acclimation (F_1,32_ = 39.93; p < 0.001) reduced citrate synthase activity while the effects of exposure to Cd (F_1,32_ = 0.93; p = 0.343) and hypoxia (F_1,32_ = 2.45; p = 0.128) were not significant (Fig. 3A). The hypoxia × Cd interaction (F_1,32_ = 8.46; p = 0.007) was significant but those of temperature × Cd (F_1,32_ = 3.43; p = 0.073), temperature × Cd (F_1,32_ = 0.89; p = 0.354) and temperature × hypoxia × Cd (F_1,32_ = 0.36; p = 0.554) were not. The overall outcome of the exposure to the stressors was that 10 ^°^C-acclimated fish mitochondria exhibited higher citrate synthase activity than the warm-acclimated irrespective of the Cd and/or hypoxia exposure status. However, the citrate synthase activities were similar in 10 ^°^C- and warm-acclimated fish exposed to hypoxia alone or jointly with Cd suggesting that these treatments antagonized the effect of warm-acclimation.

**Figure 3:**
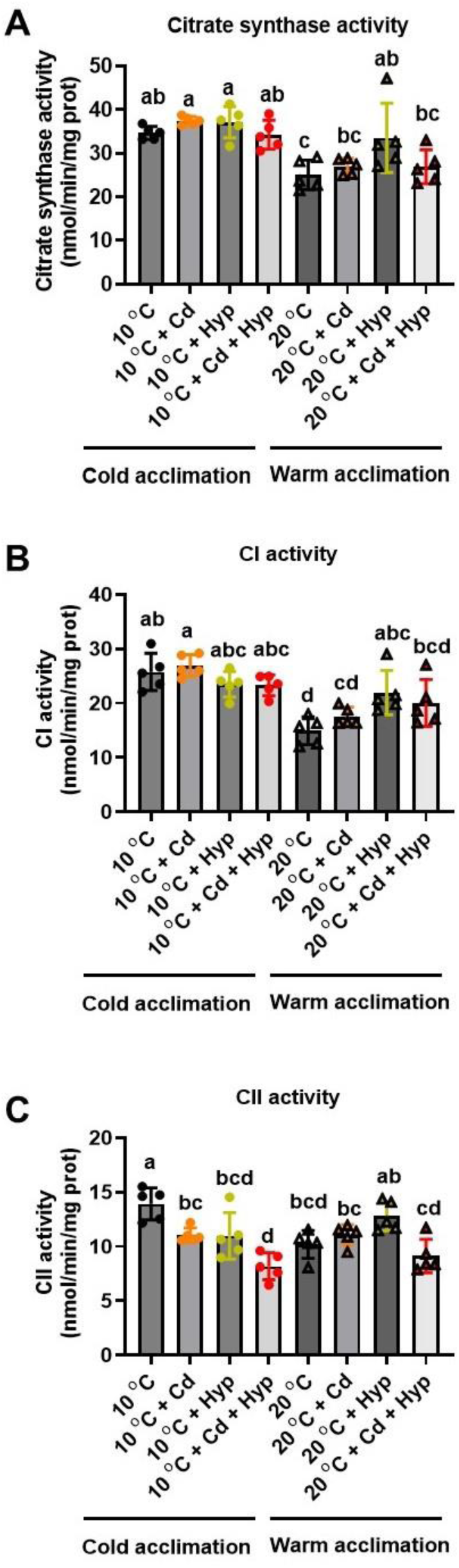
Individual and combined effects of acclimation temperature, Cd, and hypoxia on activities of mitochondrial enzymes. (**A**) citrate synthase, (**B**) complex I, and (**C**) complex II. Rainbow trout were acclimated to 10 °C (control) or 20 °C (warm-acclimated) for 50 days and exposed to (i) 10 µg/l Cd for 24 h, (ii) hypoxia (30% air saturation) for 2h, or (iii) 10 µg/l Cd for 24 h combined with hypoxia (30% air saturation) for 2 h. Liver mitochondria were isolated, and activities of the enzymes were measured. Data are means ± SEM, N = 5 independent fish. Bars with different letters represent statistically significant means (p < 0.05), three-way ANOVA, Tukey’s HSD test.

CI enzyme activity (Fig. 3B) was reduced by warm temperature acclimation (F_1,32_ = 44.99; p < 0.001) while effects of hypoxia (F_1,32_ = 0.93; p = 0.342) and Cd (F_1,32_ = 0.22; p = 0.645) were not significant. The temperature × hypoxia (F_1,32_ = 17.06; p < 0.001) interaction was significant but those of hypoxia × Cd (F_1,32_ = 2.56; p = 0.120), temperature × Cd (F_1,32_ = 0.00; p = 0.963), and temperature × hypoxia × Cd (F_1,32_ = 0.73; p = 0.400) were not significant. Pairwise comparisons of the means revealed that all the 10 ^°^C-acclimated groups exhibited higher CI enzyme activities compared with the warm-acclimated irrespective of the Cd and/or hypoxia exposure status. Interestingly, hypoxia alone and in combination with Cd mitigated the inhibitory effect of warm temperature acclimation on CI activity.

CII enzyme activity (Fig. 3C) was altered by hypoxia (F_1,32_ = 8.55; p = 0.006) and Cd (F_1,32_ = 23.23; p < 0.001) but not warm acclimation (F_1,32_ = 0.30; p = 0.586). Additionally, the interactions of hypoxia × temperature (F_1,32_ = 13.55; p = 0.001) and hypoxia × Cd were significant indicating that the effect of hypoxia on CII enzyme activity depended on temperature and Cd and vice versa. There was a significant (F_1,32_ = 6.77; p = 0.014) ternary (hypoxia × temperature × Cd) interaction suggesting that the effect of one of the factors on the CII enzyme activity was not consistent at all combinations of the other two factors. Pairwise comparisons of the means revealed that CII activity in warm-acclimated fish was significantly different only from the 10 ^°^C-acclimated group. The 10 ^°^C-acclimated fish exposed to Cd and hypoxia combined exhibited the lowest CII activity that was significantly different from all the other groups except the 10 ^°^C-acclimated exposed to hypoxia and the warm-acclimated exposed to Cd and hypoxia combined. These results indicate that combined exposure to Cd and hypoxia exacerbated the inhibitory effects of Cd on CII activity.

### 3.4. Effects of acclimation temperature, Cd, and hypoxia on CI respiratory activity plasticity: acute temperature rise

To illuminate the thermal plasticity of rainbow trout liver mitochondria, we compared the respiration rates in fish acclimated to 10 ^°^C (control) and 20 ^°^C (warm-acclimated) with those measured after acute increase in temperature (*in vitro*) from 10 to 20 ^°^C following *in vivo* exposure to Cd and/or hypoxia. For CI state 3 respiration rate (Fig. 4A), the overall group effect was significant (F_11,48_ = 15.5, p < 0.001). Specifically, acute temperature rise greatly increased state 3 respiration rate while warm acclimation had no effect. Relative to the respective temperature regimes, exposure to both Cd and hypoxia, individually and in combination, did not significantly alter state 3 respiration rates measured at the acclimation temperatures and after acute temperature rise. However, acute temperature rise induced higher state 3 respiration in mitochondria of fish exposed to hypoxia relative to those exposed to Cd alone and in combination with hypoxia.

**Figure 4:**
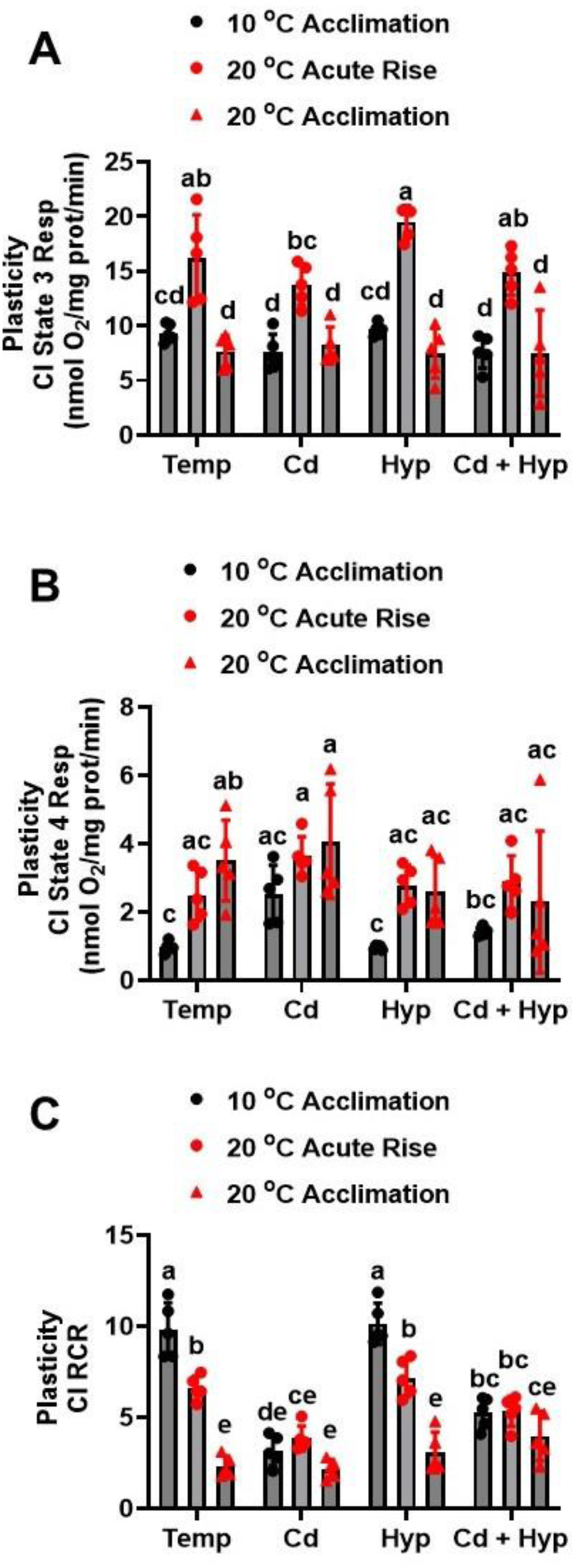
Plasticity of mitochondrial complex I powered respiration following acute temperature rise. (**A**) complex I state 3, (**B**) complex I state 4, and (**C**) complex I RCR. Rainbow trout acclimated to 10 ^°^C for 50 days were exposed to (i) 10 µg/l Cd for 24 h, (ii) hypoxia (30% air saturation) for 2h, or (iii) 10 µg/l Cd for 24 h combined with hypoxia (30% air saturation) for 2 h. Liver mitochondria were isolated and the respiration fueled by glutamate-malate was measured at 20 ^°^C. Data for the 10 ^°^C- and 20 ^°^C-acclimated fish measured at the respective acclimation temperatures were imbedded with the acute temperature rise measurements for statistical analysis. Data are means ± SEM, N = 5 independent fish. Bars with different letters represent statistically significant means (p < 0.05), three-way ANOVA, Tukey’s HSD test.

CI state 4 respiration rates (Fig. 4B) during the acute temperature rise trial were different among the experimental groups (H_(11)_ = 36.82, p <0.001). Warm-acclimated fish mitochondria and those submitted to acute temperature rise exhibited higher CI state 4 respiration rates compared with those acclimated to 10 ^°^C measured at 10 ^°^C under all conditions of exposure to hypoxia and Cd. Additionally, Cd exposure increased CI state 4 respiration rate in the 10 ^°^C- acclimated fish mitochondria measured at 10 ^°^C but the rate decreased to a level comparable to the control when Cd was combined with hypoxia.

The RCR for CI (Fig. 4C) during the acute temperature rise trial varied significantly among the treatment groups (F_11,48_ = 32.0, p < 0.001) and was highest in 10 ^°^C-acclimated fish mitochondria measured at 10 ^°^C. When 10 ^°^C-acclimated fish mitochondria were subjected to acute temperature rise, CI RCR did not change significantly; however, warm acclimation evoked a highly significant decrease in the RCR. Relative to the respective temperature acclimation group, Cd exposure reduced the RCR in mitochondria from 10 ^°^C-acclimated fish measured at 10 ^°^C and in those submitted to acute temperature rise; however, it did not change the RCR in warm-acclimated fish mitochondria measured at 20 ^°^C. Hypoxia did not alter the CI RCR relative to the respective temperature groups. However, hypoxic mitochondria acclimated to 10 ^°^C measured at 10 ^°^C or subjected to acute temperature rise had higher RCR values than the respective groups exposed to Cd. Lastly, mitochondria from fish exposed to both Cd and hypoxia exhibited similar RCR values following acute temperature rise and at respective acclimation temperatures. These RCR values were lower than for mitochondria from 10 ^°^C-acclimated fish without and with acute temperature rise under both normoxia and hypoxia, but higher than for the warm-acclimated control. Additionally, relative to the warm-acclimated control, exposure to Cd and hypoxia combined increased CI RCR in warm acclimated fish mitochondria measured at 20 ^°^C.

### 3.5. Effects of acclimation temperature, Cd, and hypoxia on CI respiratory activity plasticity: acute temperature drop

During the acute temperature drop trial, CI state 3 respiration rates (Fig. 5A) were different among the treatment groups (H_(11)_ = 40.12, p < 0.001). Warm-acclimated fish mitochondria measured at 10 ^°^C exhibited lower CI state 3 respiration compared with both the warm-acclimated measured at 20 °C and the 10 °C-acclimated measured at 10 ^°^C. Exposure to Cd and hypoxia alone or in combination did not alter CI state 3 respiration rates in mitochondria from warm- and 10 ^°^C-acclimated fish measured at the respective acclimation temperatures or the warm-acclimated after acute temperature drop.

**Figure 5:**
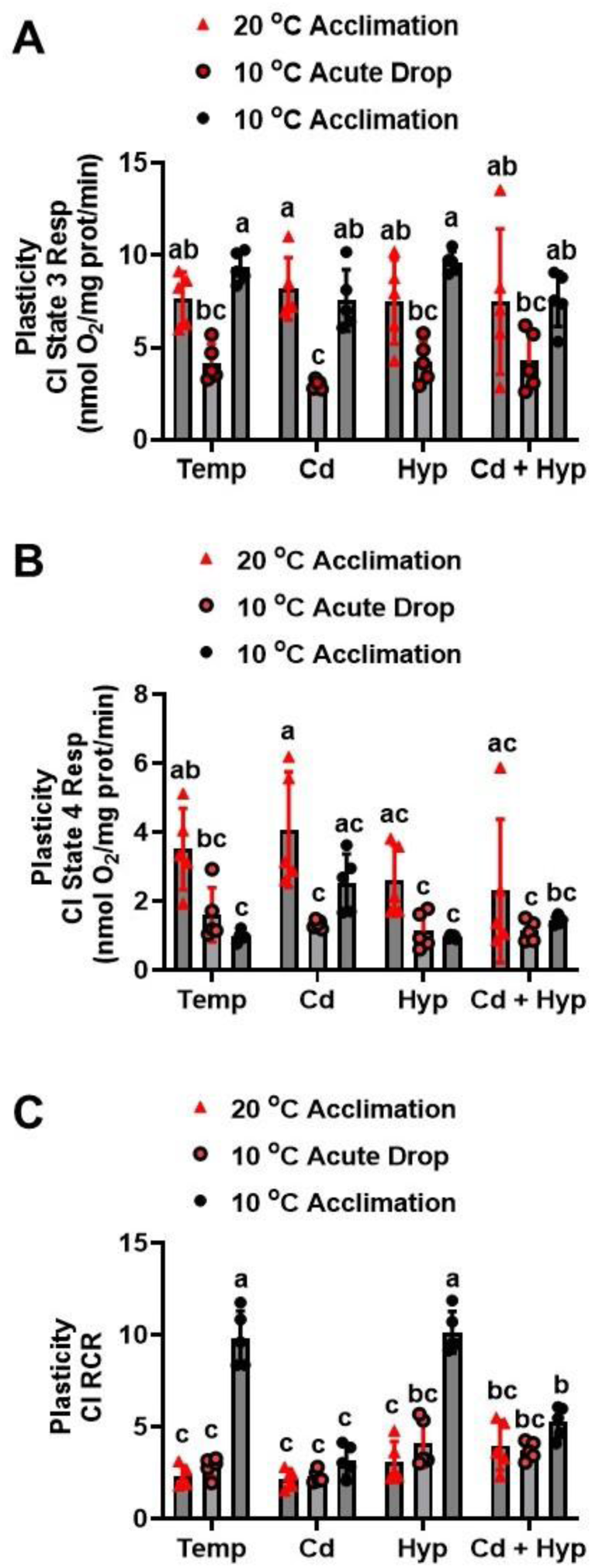
Plasticity of mitochondrial complex I powered respiration following acute temperature drop. (**A**) complex I state 3, (**B**) complex I state 4, and (**C**) complex I RCR. Rainbow trout acclimated to 20 ^°^C for 50 days were exposed to (i) 10 µg/l Cd for 24 h, (ii) hypoxia (30% air saturation) for 2h, or (iii) 10 µg/l Cd for 24 h combined with hypoxia (30% air saturation) for 2 h. Liver mitochondria were isolated and the respiration fueled by glutamate-malate was measured at 10 ^°^C. Data for the 10 ^°^C- and 20 ^°^C-acclimated fish measured at the respective acclimation temperatures were imbedded with the acute temperature drop measurements for statistical analysis. Data are means ± SEM, N = 5 independent fish. Bars with different letters represent statistically significant means (p < 0.05), three-way ANOVA, Tukey’s HSD test.

CI state 4 respiration rates (Fig. 5B) during the acute temperature drop trial were different among the treatment groups (F_11,48_ = 6.55, p < 0.001). At the respective acclimation temperatures, mitochondria from 10 ^°^C-acclimated fish exhibited lower state 4 respiration rates compared with the warm-acclimated. Acute temperature drop caused CI state 4 respiration rate to decrease to the 10 ^°^C-acclimated fish mitochondria level. Relative to the respective temperature acclimation group, Cd exposure did not alter CI state 4 respiration in warm-acclimated fish, but it increased it compared with 10 ^°^C-acclimated fish mitochondria measured at 10 ^°^C. Hypoxia alone and in combination with Cd did not significantly alter CI state 4 respiration rate relative to the respective temperature group. CI state 4 respiration rates in mitochondria from warm-and 10 ^°^C-acclimated fish exposed to hypoxia alone and in combination with Cd were lower than for the respective Cd-exposed counterparts. However, Cd combined with hypoxia did not alter CI state 4 respiration relative to the respective temperature treatment groups.

CI RCR (Fig. 5C) following acute temperature drop trial varied significantly among the treatment groups (F_48_,_11_ = 30.55, p < 0.001). Mitochondria from warm-acclimated fish measured at 20 ^°^C and after acute temperature drop had comparable RCR values which were lower than that of the 10 ^°^C-acclimated fish measured at 10 ^°^C. Relative to the respective temperature group, Cd exposure greatly reduced CI RCR in 10 ^°^C-acclimated fish mitochondria measured at 10 ^°^C without affecting it in the warm-acclimated measured at 20 °C and after acute temperature drop. The only significant effect of hypoxia exposure relative to the temperature treatment was increased CI RCR in fish submitted to acute temperature drop. For exposure to Cd and hypoxia combined, CI RCR in mitochondria from warm-acclimated fish measured at 20 ^°^C was comparable to those measured after acute temperature drop and the 10 ^°^C-acclimated measured at 10 ^°^C. Moreover, the RCR in mitochondria from fish acclimated to 10 ^°^C measured at 10 ^°^C was higher compared with the warm-acclimated after acute temperature drop. Relative to the respective temperature regimes, the CI RCR values were higher following exposure to Cd and hypoxia combined in warm-acclimated fish mitochondria measured at 20 ^°^C but lower in 10 ^°^C- acclimated fish mitochondria measured at 10 ^°^C.

### 3.6. Effects of acclimation temperature, Cd, and hypoxia on CI + II respiratory activity plasticity: acute temperature rise

For CI + II state 3 respiration response to acute temperature rise (Fig. 6A), the group effect was significant (F_11,48_ = 28.0, p < 0.001). Acute temperature rise greatly increased CI + II state 3 respiration rate and this increase was blunted by exposure to Cd alone and in combination with hypoxia. In addition, exposure to Cd alone and jointly with hypoxia but not hypoxia alone reduced state 3 respiration rate in mitochondria from fish acclimated to 10 ^°^C measured at 10 ^°^C.

**Figure 6:**
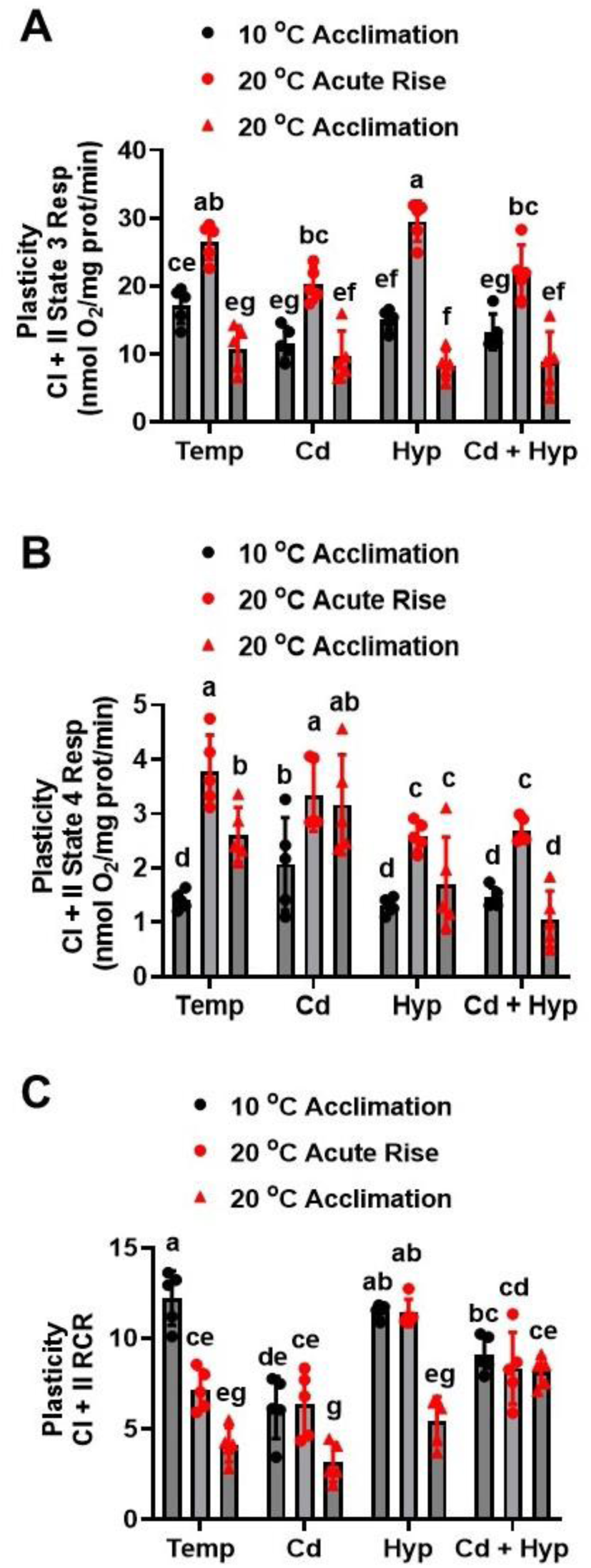
Plasticity of mitochondrial complex I + II powered respiration following acute temperature rise. (**A**) state 3, (**B**) state 4, and (**C**) RCR. Rainbow trout acclimated to 10 ^°^C for 50 days were exposed to (i) 10 µg/l Cd for 24 h, (ii) hypoxia (30% air saturation) for 2h, or (iii) 10 µg/l Cd for 24 h combined with hypoxia (30% air saturation) for 2 h. Liver mitochondria were isolated and the respiration fueled by glutamate-malate-succinate was measured at 20 ^°^C. Data for the 10 ^°^C- and 20 ^°^C-acclimated fish measured at the respective acclimation temperatures were imbedded with the acute temperature rise measurements for statistical analysis. Data are means ± SEM, N = 5 independent fish. Bars with different letters represent statistically significant means (p < 0.05), three-way ANOVA, Tukey’s HSD test.

CI + II state 4 respiration rates (Fig. 6B) changed significantly among the experimental groups (H_(11)_ = 42.7, p <0.001) during the acute temperature rise trial. Mitochondria from 10 ^°^C- acclimated fish submitted to acute temperature rise exhibited higher CI + II state 4 respiration rate compared with those acclimated to 10 ^°^C measured at 10 ^°^C under all conditions. Warm-acclimation alone and in combination with Cd increased CI + II state 4 respiration while hypoxia alone and in combination with Cd exposure blunted the stimulatory effect of warm acclimation and acute temperature rise. Similarly, CI + II state 4+ (Fig. S2A) varied significantly among the treatment groups (F_11,48_ = 22.8, p < 0.001) and mirrored the CI + II state 4 response pattern.

CI+II RCR during the acute temperature rise trial (Fig. 6C) varied among the treatment groups (F_11,48_ = 25.9, p < 0.001), being highest in 10 ^°^C-acclimated fish mitochondria measured at 10 ^°^C and decreasing upon acute temperature rise and warm acclimation. Cd exposure reduced CI + II RCR in 10 ^°^C-acclimated fish mitochondria measured at 10 ^°^C but did not alter it in mitochondria subjected to acute temperature rise or warm acclimation. Hypoxia increased CI + II RCR in mitochondria subjected to acute temperature rise without altering it in 10 ^°^C- and warm-acclimated fish mitochondria measured at the respective acclimation temperatures. In contrast, Cd combined with hypoxia reduced RCR in 10 ^°^C-acclimated fish mitochondria measured at 10 ^°^C and increased it in warm-acclimated fish mitochondria measured at 20 ^°^C. The CI + II RCR+ response to acute temperature rise (Fig. S2B) mirrored that of CI + II RCR and differed significantly among the treatment groups (F_11,48_ = 14.4, p < 0.001).

### 3.7. Effects of acclimation temperature, Cd, and hypoxia on CI + II respiratory activity plasticity: acute temperature drop

In the acute temperature drop test, CI + II state 3 respiration rates (Fig. 7A) were different among the treatment groups (F_11,48_ = 12.7, p < 0.001). Warm-acclimated fish mitochondria subjected to acute temperature drop exhibited lower CI + II state 3 respiration rate compared with 10 ^°^C-acclimated fish mitochondria measured at 10 ^°^C under all the experimental conditions. In contrast, except for the Cd-exposed group, all the CI + II state 3 respiration rates measured after acute temperature drop were comparable to those of the warm-acclimated fish mitochondria measured at 20 ^°^C. Additionally, exposure to Cd but not hypoxia or Cd + hypoxia reduced state 3 respiration rate after acute temperature drop relative to the 10 ^°^C-acclimated fish mitochondria measured at 10 ^°^C.

**Figure 7:**
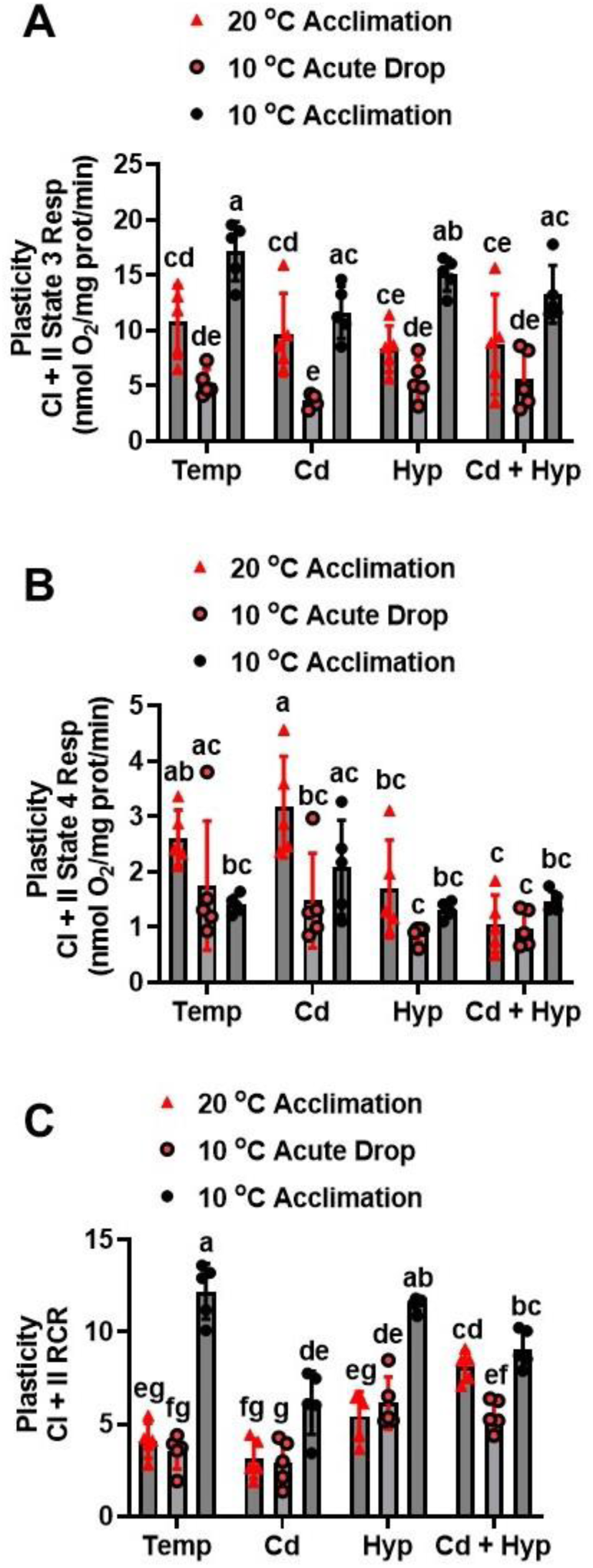
Plasticity of mitochondrial complex I + II powered respiration following acute temperature drop. (**A**) state 3, (**B**) state 4, and (**C**) RCR. Rainbow trout acclimated to 20 ^°^C for 50 days were exposed to (i) 10 µg/l Cd for 24 h, (ii) hypoxia (30% air saturation) for 2h, or (iii) 10 µg/l Cd for 24 h combined with hypoxia (30% air saturation) for 2 h. Liver mitochondria were isolated and the respiration fueled by glutamate-malate-succinate was measured at 10 ^°^C. Data for the 10 ^°^C- and 20 ^°^C-acclimated fish measured at the respective acclimation temperatures were imbedded with the acute temperature rise measurements for statistical analysis. Data are means ± SEM, N = 5 independent fish. Bars with different letters represent statistically significant means (p < 0.05), three-way ANOVA, Tukey’s HSD test.

CI + II state 4 respiration rates (Fig. 7B) varied among the treatment groups (H_11,48_ = 5.7, p < 0.001) during the acute temperature drop trial. The temperature regime did not significantly alter CI + II state 4 respiration rate and exposure to Cd or hypoxia did not change the effect of temperature in corresponding groups. However, warm-acclimated fish mitochondria exposed to Cd had higher CI + II state 4 respiration compared with mitochondria subjected to acute temperature drop as well as all the groups exposed to hypoxia alone and in combination with Cd. Exposure to hypoxia alone or in combination with Cd in warm acclimated fish mitochondria measured at 20 ^°^C decreased CI + II state 4 respiration rate thereby abolishing the difference relative to the 10 ^°^C-acclimated measured at 10 ^°^C. Relative to the temperature regime, CI + II state 4 respiration rate in mitochondria subjected to acute temperature drop was not altered by exposure to Cd or hypoxia singly and in combination. CI + II state 4+ respiration rate during the acute temperature drop trial (Fig. S3A) varied among the treatment groups (F_11,48_ = 5.9, p < 0.001), with a pattern comparable to that of RCR.

CI + II RCR for the acute temperature drop trial (Fig. 7C) varied significantly among the treatment groups (F_11,48_ = 37.1, p < 0.001). Mitochondria from warm-acclimated fish measured at 20 ^°^C and after acute temperature drop had comparable CI + II RCR values which were substantially lower than for the 10 ^°^C-acclimated measured at 10 ^°^C. Cd exposure markedly reduced CI + II RCR in 10 ^°^C-acclimated fish mitochondria measured at 10 ^°^C without affecting it in warm-acclimated fish measured at 20 ^°^C and after acute temperature drop. Following exposure to hypoxia, mitochondria of warm-acclimated fish measured at either 20 ^°^C or 10 ^°^C had comparable CI + II RCR values which were lower than for hypoxic 10 ^°^C-acclimated fish mitochondria measured at 10 ^°^C. Additionally, CI + II RCR for mitochondria from fish exposed to hypoxia and subjected to acute temperature drop were higher compared with the respective normoxic temperature regime counterpart. For the combined Cd and hypoxia exposure, CI + II RCR in mitochondria from warm-acclimated fish measured at 20 ^°^C was comparable to that of the 10 ^°^C-acclimated measured at 10 ^°^C but significantly higher than for mitochondria subjected to acute temperature drop. Compared with the respective temperature group, exposure to Cd and hypoxia jointly increased CI + II RCR in warm-acclimated mitochondria measured at 20 ^°^C and those subjected to acute temperature drop; however, the RCR in 10 ^°^C-acclimated fish mitochondria exposed to Cd and hypoxia jointly measured at 10 ^°^C was reduced. CI + II RCR+ for the acute temperature drop trial (Fig. S3B) varied significantly among the treatment groups (F_11,48_ = 19.1, p < 0.001), with a pattern comparable to that of RCR.

## 4.0. Discussion

Fish have evolved elaborate biochemical, physiological, and behavioral mechanisms that enable them to survive and function in their natural habitats. These mechanisms are challenged by climate change that is not only predicted to cause gradual overall warming of water bodies but increase the frequency and intensity of temperature fluctuations (heatwaves) as well as extreme hypoxic events (Frölicher et al., 2018; Oliver et al., 2018; Eahart et al., 2022; Harvey et al., 2022). Moreover, temperature changes occur together with other stress factors concurrently or sequentially and interact both at the environmental and organism levels. Our study probed the effects of warm acclimation, hypoxia, and Cd exposure on mitochondrial bioenergetics and how they affect mitochondrial functional plasticity. We found pervasive effects of warm acclimation on mitochondrial bioenergetics that were moderated by more subtle effects of Cd and hypoxia. Specifically, hypoxia and/or Cd at the levels and extents tested either mitigated or exacerbated the effects of warm acclimation.

### 4.1. Effects and interactions of temperature, Cd, and hypoxia on mitochondrial bioenergetics

*In vitro* studies with isolated mitochondria are important in elucidating mechanisms of action of stressors but are devoid of moderating adaptive responses that occur following exposure to stressors *in vivo*. We uncovered notable similarities and differences between reported *in vitro* observations using isolated organelles and our *in vivo* study. Warm acclimation decreased state 3 respiration driven by CI + II but not CI. These observations on the effect of warm acclimation contrast the reported stimulatory effect of acute temperature rise on state 3 respiration (Zukiene et al., 2010; Galli and Richards, 2012; Rodnick et al., 2014; Sappal et al., 2014; Onukwufor et al., 2016). Stimulation of state 3 respiration by acute temperature rise is attributed to pervasive increased rates of biological/physiological processes with temperature and altered membrane fluidity and composition (Dahlhoff and Somero, 1993; Kraffe et al., 2007; Chung et al., 2018). Time-course assessments of effect of temperature change revealed a biphasic response to warm acclimation in which mitochondrial oxidative capacity initially increase and then decrease to the level prior to temperature rise (Bouchard and Guderley, 2003; Kraffe et al., 2007). In agreement with this theme, state 3 respiration rates for both CI and CI + II in 10 °C-acclimated and warm (20 °C)-acclimated fish mitochondria were similar, indicating reversal of initial acute temperature induced stimulatory effect (Fig. 4; Fig. 6).

Our observation that the effects of warm acclimation on CI and CI + II driven state 3 respirations were similar suggests a common underlying mechanism such as altered membrane composition and fluidity (Kraffe et al., 2007) and contrast previous studies that found CI respiration was enhanced by warm acclimation (Gerber et al., 2020, 2021). Moreover, our findings are inconsistent with the proposal that CI is the primary driver of mitochondrial bioenergetic effects of warm acclimation (Pichaud et al., 2017; Strobel et al., 2013; Gerber et al., 2020, 2021) because CI + CII driven respiration was reduced. While it could be argued that the reduced CI + II respiration was due to reduction of the CI component because we did not measure CII driven respiration alone, reduced catalytic activity of CII in warm-acclimated fish mitochondria (Fig. 3C) supports inhibition of CII driven respiration. Regardless, differences in fish species, tissue/organ, and temperature regimes employed among studies can account for some of the reported discrepancies in mitochondrial functional responses following warm acclimation.

Cd exposure inhibited state 3 respiration driven by CI + CII but not CI contrasting previous reports of inhibition of CI-driven respiration under *in vitro* conditions (Sokolova 2004, Wang et al., 2004, Ivanina et al., 2011; Onukwufor et al., 2017). Our study suggests that maximally activated ETS (CI + CII respiration) is more responsive to warm acclimation and Cd than partially activated ETS (CI driven respiration) and would be a more sensitive endpoint for assessing effects and interactions of stressors on mitochondrial bioenergetics. Unlike warm acclimation and Cd, *in vivo* exposure of rainbow trout to hypoxia for 2 h did not impact CI and CI + CII driven respirations contrasting previous studies that found inhibition of state 3 respiration by acute hypoxia exposure in hypoxia-sensitive species (Navet et al., 2006; Hickey et al., 2012; Onukwufor et al., 2014, 2016, 2017). Moreover, in our previous study (Onukwufor et al., 2016), exposure to hypoxia combined with acute temperature stress exacerbated the inhibitory effect on CI driven state 3 respiration relative to the stressors singly. In our present *in vivo* study, co-exposure to warm acclimation and hypoxia did not lead to added response beyond that observed with warm temperature acclimation.

What are the possible reasons why most of the commonly observed acute responses on state 3 respiration *in vitro* were not replicated in an *in vivo* setting? A potential explanation is that the intracellular O_2_ tension achieved under the hypoxia regime used was not low enough to alter mitochondrial bioenergetic activity. Indeed, the fish were exposed to 30% air saturation for only 2 h, while isolated mitochondria in our earlier *in vitro* studies were exposed to severe hypoxia (0.1 mg O_2_/l) bordering on anoxia (Onukwufor et al., 2016, 2017). It could also be that under the *in vivo* exposure conditions, fish recruited adaptive (biochemical, physiological, and behavioral) mechanisms to counteract the deleterious effects of hypoxia (Richards, 2011), while under *in vitro* conditions isolated mitochondrial lack protective adaptive mechanisms.

Warm acclimation stimulated CI and CI + II powered leak respiration akin to several previous *in vitro* studies in which isolated mitochondria were subjected to acute temperature rise (Onukwufor et al., 2015, Sappal et al., 2015; 2016). This implies that increased membrane leakiness commonly observed after acute temperature rise (Kraffe et al., 2007; Pichaud et al., 2017) persists upon warm acclimation. Similarly, Cd exacerbated the effect of warm acclimation on leak respiration powered by both CI and CI + II in agreement with *in vitro* observations (Adiele et al., 2012; Onukwufor et al., 2016). In contrast, exposure to hypoxia *in vivo* with and without warm acclimation did not alter leak respiration powered by both CI and CI + II. These findings contrast those of *in vitro* exposure of isolated mitochondria to hypoxia-reoxygenation in which leak respiration was stimulated (Navet et al., 2006; Hickey et al., 2012; Onukwufor et al., 2016), with the authors attributing the effect to increased membrane leakiness. Although the fish in our *in vivo* study were removed from the hypoxic water and immediately sacrificed for liver mitochondrial isolation, the mitochondria did get re-oxygenated during isolation and measurement of respiration. Thus, the combination of *in vivo* hypoxia and *in vitro* re-oxygenation did not produce the same results compared with when both procedures were done *in vitro* with isolated mitochondria.

Several studies have shown that acute exposure of isolated mitochondria to warm temperature causes a significant reduction in RCR (Onukwufor et al., 2015; 2017, Sappal et al., 2015, Chung and Schulte, 2015; Chung et al., 2017). Consistent with these studies, warm acclimation reduced RCR for both CI and CI + II powered respirations. Similarly, *in vivo* Cd exposure reduced RCR for respirations driven by both CI and CI + II in agreement with reports that impairment of mitochondrial bioenergetic function is a hallmark of Cd exposure (Sokolova, 2004; Onukwufor et al., 2015). This reduction of phosphorylation efficiency (RCR) could be in part due to the inhibition of enzymes of the ETS and/or citric acid cycle (CAC) (Onukwufor et al., 2014, Belyaeva and Korotkov, 2003). Indeed, we show that Cd exposure reduced the activity of CII (Fig. 3). The reduction of RCR by Cd was exacerbated by warm acclimation in agreement with previous reports of effects of combined exposure to Cd and acute temperature rise in isolated mitochondria (Sokolova 2004, Onukwufor et al., 2015; 2017). Thus, the impact of Cd on mitochondrial respiratory function is similar under both *in vitro* and *in vivo* exposure regimes.

Acute hypoxia-reoxygenation exposure reduced RCR in isolated mitochondria, an effect that was exacerbated by warm temperature stress (Navet et al., 2006; Hickey et al., 2012; Onukwufor et al., 2016). We found different effects under *in vivo* conditions wherein hypoxia did not alter CI RCR while CI + II RCR was marginally reduced. The RCR for both CI and CI + II powered respirations were higher for the ternary combination of the stressors compared with the individual and binary effects of Cd and warm acclimation indicating that the ameliorative effects of *in vivo* exposure to the three stressors combined was driven by hypoxia. This suggests that hypoxia exposure elicited adaptive stress response mechanisms that ameliorated the effects of warm acclimation and/or Cd on mitochondrial function *in vivo*. In this regard, chronic hypoxia was reported to alter cardiac mitochondrial membrane composition in the sablefish, *Anaplopoma fimbria* (Gerber et al., 2019). The duration of hypoxia in our study was likely not long enough to cause changes in membrane composition. Thus, future studies should probe potential mechanisms by which a short hypoxic event *in vivo* protects mitochondria against warm temperature stress and/or Cd toxicity.

### 4.2. Effects of temperature, Cd, and hypoxia on activities of mitochondrial enzymes

The activities of citrate synthase, CI, and CII were all reduced by warm acclimation in agreement with previous studies in a variety of organisms that reported higher mitochondrial enzyme activities at cooler relative to warmer temperatures (Bouchard and Guderley, 2003; Guderley, 2004; Kraffe et al., 2007; Fangue et al., 2009; Hoffschröer et al., 2024). Reduced citrate synthase activity in warm acclimated organisms is thought to result from quantitative changes mediated by gene expression (Somero et al., 2017). Because citrate synthase activity is an indicator of substrate flux in CAC and mitochondrial density (Moyes et al.,1997; Lucassen et al., 2003; Hoffschröer et al., 2024), its higher activity in mitochondria of fish acclimated to 10 ^°^C relative to 20 ^°^C suggests enhanced substrate oxidation with increased delivery of reduced equivalents to the ETS. Additionally, the higher CI and CII activities in the 10 ^°^C-acclimated fish compared with the warm-acclimated indicate that flux through the ETS subsystem was reduced by warm acclimation. Therefore, in our study, warm acclimation reduced mitochondrial oxidative capacity by downregulating both the substrate oxidation (CAC) and the ETS subsystems.

The effects of Cd and hypoxia on mitochondrial enzyme activities were less pronounced and more variable compared with those of warm acclimation. Cd and hypoxia did not alter activities of citrate synthase and CI but both factors reduced CII activity and the effect of Cd depended on DO status. Surprisingly, while hypoxia alone and jointly with Cd mitigated the inhibitory effect of warm temperature acclimation on citrate synthase activity, hypoxia exacerbated the inhibitory effect of Cd on CII activity. Overall, our study shows that the interactive effects of multiple stressors vary with the context.

### 4.3. Plasticity of CI and CI + II respiratory functions

Mitochondria are plastic organelles that undergo morphological and functional changes in response to cellular signals and environmental stress. However, knowledge of the effect of warm acclimation with acute thermal shifts and exposure to metals and hypoxia on mitochondrial respiration is lacking. Concurrent investigation of effects of warm acclimation and acute temperature shifts offers the opportunity to unveil and compare the stable and transient mitochondrial responses to thermal stress. Here, we compared the effects of long-term acclimation to 10 ^°^C and 20 ^°^C *in vivo* with those of *in vitro* acute temperature shift (rise: 10 ^°^C → 20 ^°^C and drop: 20 ^°^C → 10^°^C).

Acute temperature rise increased both state 3 and leak respirations but decreased the RCR because of preferential increase in the latter. This response has been demonstrated in several studies involving acute temperature stress (Onukwufor et al., 2015; 2017; Sappal et al., 2015, Chung and Schulte, 2015; Chung et al., 2017). Upon acclimation to 20 ^°^C the pattern changed with state 3 respiration returning to the 10 ^°^C-acclimation (control) level while leak respiration increased above the acute phase rate. Thus, a key adverse effect of warm temperature acclimation on mitochondrial respiration is the inability of the leak respiration to return to the pre-temperature rise level. This indicates that while acclimation to warm temperature results in overall compensation of the acute temperature rise induced increase in O_2_ consumption, the proportion of leak respiration increased.

For acute temperature drop, among the three groups, the 20-^°^C acclimated fish mitochondria exhibited intermediate state 3 respiration and highest leak respiration rates both of which decreased markedly upon acute temperature drop to 10 ^°^C. The decrease in state 3 respiration was much greater relative to that of the state 4 respiration resulting in a low RCR indicative of poor mitochondrial coupling and low oxidative phosphorylation capacity. In contrast, 10 ^°^C-acclimated fish mitochondria had the highest state 3 respiration and lowest leak respiration, and consequently the highest RCR. This suggests that reducing leakiness of the mitochondrial membranes improves electron flux in the ETS subsystem thus increasing the oxidative phosphorylation capacity.

## Conclusions

While our study revealed pervasive inhibitory and stimulatory effects of warm acclimation on mitochondrial bioenergetic states, the key effect of warm acclimation was reduced oxidative phosphorylation efficiency due to sustained preferential stimulation of the proton leak subsystem. Specifically, respiration linked to phosphorylation (state 3) returned to pre-temperature rise level after acclimation, but leak respiration remained highly elevated. The dominance of the effect of warm acclimation compared with those of Cd and hypoxia in our study likely reflects the experimental regime, wherein the levels and durations of Cd and hypoxia exposures were lower compared with those of warm acclimation. Overall, our study indicates that joint actions of temperature, Cd, and hypoxia on mitochondrial bioenergetics cannot be deduced from their individual effects.

## Supporting information

Supplemental figure 1-3, and will be used for the link to the file on the preprint site

## Data availability

All data associated with the study will be made available upon request.

