## Supplemental figure 1-3, and will be used for the link to the file on the preprint site for "Interactive effects of temperature, cadmium, and hypoxia on rainbow trout *(Oncorhynchus mykiss)* liver mitochondrial bioenergetics"

**S1**

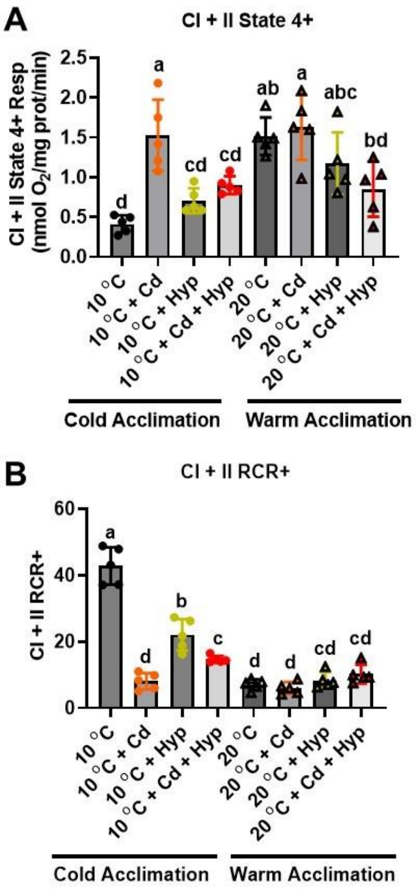


**S1**: Individual and combined effects of acclimation temperature, Cd, and hypoxia on mitochondrial complex I + II powered respiration. (**A**) state 4+, and (**B**) RCR+. Rainbow trout were acclimated to 10 °C (control) or 20 °C (warm-acclimated) for 50 days and exposed to (i) 10 µg/l Cd for 24 h, (ii) hypoxia (30% air saturation) for 2h, or (iii) 10 µg/l Cd for 24 h combined with hypoxia (30% air saturation) for 2 h. Liver mitochondria were isolated and the respiration fueled by glutamate-malate-succinate was measured at the respective acclimation temperature. Data are means ± SEM, N = 5 independent fish. Bars with different letters represent statistically significant means ($p < 0.05$), three-way ANOVA, Tukey's HSD test.

**S2**

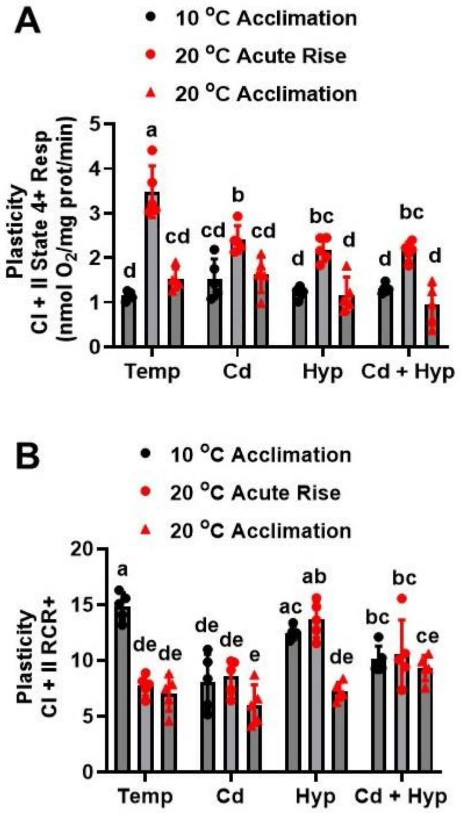


**S2**: Plasticity of mitochondrial complex I + II powered respiration following acute temperature rise. (**A**) state 4+, and (**B**) RCR+. Rainbow trout acclimated to 10 ˚C for 50 days were exposed to (i) 10 µg/l Cd for 24 h, (ii) hypoxia (30% air saturation) for 2h, or (iii) 10 µg/l Cd for 24 h combined with hypoxia (30% air saturation) for 2 h. Liver mitochondria were isolated and the respiration fueled by glutamate-malate-succinate was measured at 20 ˚C. Data for the 10 ˚C- and 20 ˚C-acclimated fish measured at the respective acclimation temperatures were imbedded with the acute temperature rise measurements for statistical analysis. Data are means ± SEM, N = 5 independent fish. Bars with different letters represent statistically significant means ($p < 0.05$), three-way ANOVA, Tukey's HSD test.

**S3**

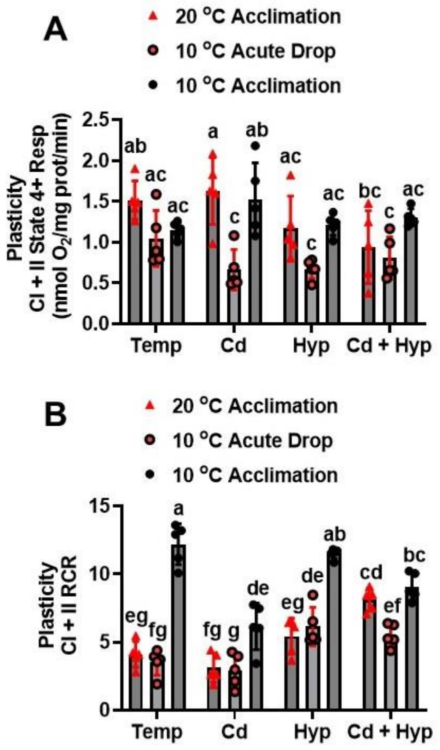


**S3**: Plasticity of mitochondrial complex I + II powered respiration following acute temperature drop. (**A**) state 4+, and (**B**) RCR+. Rainbow trout acclimated to 20 ˚C for 50 days were exposed to (i) 10 µg/l Cd for 24 h, (ii) hypoxia (30% air saturation) for 2h, or (iii) 10 µg/l Cd for 24 h combined with hypoxia (30% air saturation) for 2 h. Liver mitochondria were isolated and the respiration fueled by glutamate-malate-succinate was measured at 10 ˚C. Data for the 10 ˚C- and 20 ˚C-acclimated fish measured at the respective acclimation temperatures were imbedded with the acute temperature rise measurements for statistical analysis. Data are means ± SEM, N = 5 independent fish. Bars with different letters represent statistically significant means ($p < 0.05$), three-way ANOVA, Tukey's HSD test.
